# C1q Enables Influenza HA Stem Binding Antibodies to Block Viral Attachment and Broadens the Antibody Escape Repertoire

**DOI:** 10.1101/2023.06.12.544648

**Authors:** Ivan Kosik, Jefferson Da Silva Santos, Matthew Angel, Zhe Hu, Jaroslav Holly, James S. Gibbs, Tanner Gill, Martina Kosikova, Tiansheng Li, William Bakhache, Patrick T. Dolan, Hang Xie, Sarah F. Andrews, Rebecca A. Gillespie, Masaru Kanekiyo, Adrian B. McDermott, Theodore C. Pierson, Jonathan W. Yewdell

## Abstract

Broadly neutralizing, anti-hemagglutinin stem antibodies (Abs) are a promising universal influenza vaccine target. While anti-stem Abs are not believed to block viral attachment, we show that C1q confers attachment inhibition and boosts fusion and neuraminidase inhibition, greatly enhancing virus neutralization activity in vitro and in mice challenged with influenza virus via the respiratory route. These effects reflect increased steric interference and not increased Ab avidity. Remarkably, C1q greatly expands the anti-stem Ab viral escape repertoire to include residues throughout the hemagglutinin. Some substitutions cause antigenic alterations in the globular region or modulate HA receptor avidity. We also show that C1q enhances the neutralization activity of non-RBD anti-SARS-CoV-2 Spike Abs, an effect dependent on Spike density on the virion surface. Together, our findings show that first, Ab function must be considered in a physiological context and second, inferring the exact selection pressure for Ab-driven viral evolution is risky business, at best.

## Main

In the wake of COVID-19, influenza A virus (IAV) is re-exacting its annual economic and health burden. The hemagglutinin (HA) and neuraminidase (NA) glycoproteins decorating the virion and infected cell surface are the primary targets of protective antibodies (Abs)^1^. Sixteen HA serological subtypes form two evolutionary clades^2^, though only three subtypes (H1, H2, H3) are known to transmit efficiently in humans. Most anti-HA Abs bind to the region of five canonical antigenic sites in the globular head domain (Sa, Sb, Ca1, Ca2, Cb for H1 HA, A, B, C, D, E for H3 HA)^3, 4^ (Fig. 1A). The most potent head-specific Abs sterically interfere with the receptor binding site (RBS)^3, 5^. The effectiveness of anti-head Abs wanes as HA accumulates amino acid substitutions during viral evolution in humans ^3, 6^. The far more conserved HA stem is also recognized by Abs, which neutralize *in vitro*^7^ and protect *in vivo*^8^ without inhibiting viral attachment to target cells. Stem-specific Abs block viral replication by blocking HA-mediated fusion and NA-mediated release ^9, 10^. The stem evolves more slowly than the head, probably due to a combination of fitness constraints and lower selection pressure from anti-stem Abs^11–16^.

The complement system is a central player in innate and adaptive immunity. Complement proteins cascadingly collaborate to opsonize pathogens and induce inflammatory responses. C1q is an abundant serum complement protein (70 μg/ml) comprising 6 subunits that bind individual IgG Fc domains (K_D_ = 100 μM)^17^. When such Abs are appropriately arrayed on a surface, *e.g.*, virus or cell, multivalent binding increases C1 avidity enormously (K_D_ =10nM) ^18^, leading to essentially irreversible binding.

C1q is known to increase the functionality of anti-viral Abs ^19–22^, though its effects on Ab-driven viral evolution have yet to be examined. For HA, Gerhard and colleagues reported that C1q enhances the ability of anti-HA head Abs to inhibit attachment and viral infectivity^23^. Here, we extend these findings to anti-stem HA Abs and Abs specific for the S2 domain of the SARS-CoV2 Spike protein and explore the effects of C1q on Ab-driven HA evolution.

## Results

### Anti-stem Abs inhibit IAV attachment in the presence of C1q

We^10^ (and many other labs^7, 24^) reported that anti-stem Abs do not inhibit attachment of clade 1 A/Puerto Rico/8/1934 (H1N1) (PR8) or clade 2 A/Hong Kong/1968 (H3N2) (HK68) viruses. Based on findings that C1q enhances hemagglutination inhibition (HI) and virus neutralization (VN) activities of anti-HA head Abs^23^, might C1q have a similar effect on anti-stem Abs? To this end, we tested the impact of C1q on Ab attachment inhibition (AI), as measured by virus attachment to plate-bound bovine fetuin (a highly sialylated serum protein), using viral NA activity to monitor virus binding as described^25^ (Fig. 1B).

Adding C1q at a physiological concentration (70ug/ml^-1^ (160nM)) to the human anti-stem mAb 310-16G8 conferred viral attachment inhibition to the mAb, which now exhibited an AI_50_ of 14.56 nM against PR8. Using mouse mAbs specific for Sb (H28-D14) or Cb (H35-C12) sites revealed an inverse correlation between epitope distance from the RBS and C1q enhancement (stem (>1000-fold), Cb (12-fold), Sb (2.6-fold)). We next examined C1q AI enhancement of a panel of 12 anti-clade 1 or clade 2 stem mAbs (Fig. 1C). As expected, none of the mAbs alone exhibit measurable AI at a saturating concentration (200nM). Adding C1q conferred AI activity with AI_50_s ranging from 10.3nM-158nM for clade 1 and 29nM-103nM for clade 2-specific mAbs. To determine the C1q concentration required for AI enhancement, we serially diluted C1q with Abs at a saturating concentration (Supplemental Fig. 1A). Five to ten-fold lower than physiological C1q concentration was sufficient to inhibit attachment by 50% of PR8 for the five anti-stem Abs tested.

To extend these findings to serum Abs, we heat-inactivated C1q in nine adult human plasma samples. We established the presence of anti-stem Abs by testing the capacity of sera to bind recombinant stem^26, 27^ or block binding of the Fi6 stem-specific mAb via biolayer interferometry (BLI) (Fig. 1D). All but one plasma had detectable Ab that bound stem and blocked binding of Fi6 to stem. We next tested C1q’s ability to endow AI to plasma stem binding Abs. To exclude the contribution of HA head-specific antibodies present in the plasma, we used a virus with a H6 head domain genetically appended to a H1 stalk. In the absence of C1q, none of the plasma exhibit AI activity. C1q confers AI activity to all stem Ab-containing plasma but not the anti-stem Ab-negative plasma, which is an ideal negative control (Fig. 1E).

To confirm the molecular basis for the observed effects of C1q on mAb activity, we genetically introduced the Fc-K322A substitution, which diminishes C1q binding^28^, into several mAbs. With negligible effect on ELISA-based Ab apparent binding affinity measurements (Supplemental Fig. 1E), K322A almost completely abolished C1q AI activity and-VN enhancement (Fig. 1F) (to be described below) for the 310-16G8 anti-stem mAb and other stem Abs tested (Fig. 1G).

**Figure 1.**
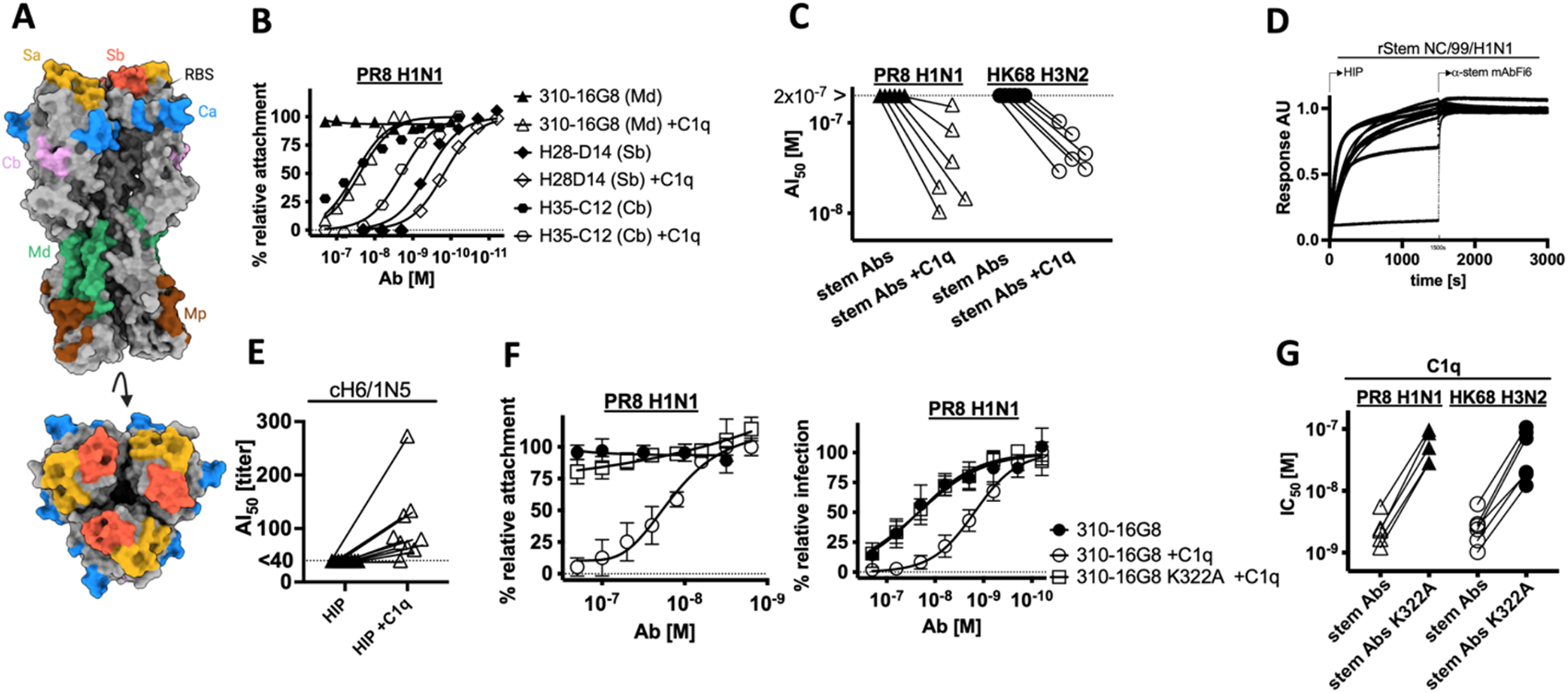
C1q confers and expands anti-HA stem Ab anti-viral functions. **(A)** PR8 hemagglutinin (PDB:1RU7), side and top views. Canonical antigenic sites are identified by color: Sa yellow, Sb orange, Ca blue, Cb pink, Md (stem membrane distal) green, and Mp (stem membrane proximal) brown. **(B)** AI activity of the indicated anti-stem and head Abs in the absence and presence of C1q *vs*. PR8 determined using the fetuin-attachment assay. **(C)** AI_50_ (concentration required to inhibit attachment of 50% of the input viral particles) for a panel of ten anti-stem Abs (8 clade-specific and two pan clade Abs) in the fetuin-attachment assay against PR8 or HK. **(D)** Anti-stem Abs in human plasma detected by BLI using recombinant stem (NC/99/H1N1) as antigen. Right side of graph shows lack of increased signal following the addition of anti-stem mAb Fi6 due to competition from plasma Abs. Note the large increase in signal in the sole plasma sample lacking anti-stem Abs (red arrow) **(E)** AI_50_ of human plasma in the fetuin-attachment assay against HA chimeric virus (H6 head/H1 stem) with and without C1q . Each experiment shows the mean of four-six technical replicates acquired in two-three independent experiments for each plasma tested. **(F)** AI and VN activity of representative BN-HA stem mAb 310-16G8 and its ΔC1q-binding variant (Fc mutation K322A) in the presence and absence of the C1q in fetuin-attachment (left) and flow cytometry single cycle VN (right) assays. **(G)** IC_50_ VN titers for a panel of 11 anti-stem Abs and K322A variants with and without C1q against viruses indicated. Each experiment shows the mean of four-six technical replicates acquired in two-three independent experiments for each plasma tested. Non-virus control (NVC) and no Ab control (NoAbC) were included in each experiment. Data are normalized using NoAbC set to 0% and plateau response observed for at least two highest Ab dilutions employed set to 100%. BLI data normalized by maximal signal acquired for individual samples. Data are plotted with GraphPad Prism7 software to generate nonlinear regression curves and calculate AI_50_ using the normalized dose-response inhibition model.

Based on these findings, we conclude that C1q confers AI activity to monoclonal and polyclonal anti-stem Abs.

### Both AI and FI contribute to VN enhancement mediated by C1q

How does C1q-enhanced AI affect VN activity? We used H1 (PR8) or H3 (HK) expressing recombinant reporter viruses that express mCherry encoded by gene segment 8 in a flow cytometry-based VN assay. By assessing early infection (5 HPI), we could exclusively measure Ab-mediated entry blockade. For a representative anti-stem mAb (310-16F8), C1q enhances VN activity ∼30-fold (Fig. 2A). All clade I and 2 anti-stem mAbs demonstrated similar VN enhancement (Fig. 2B). For human plasma, C1q enhanced VN of chimeric H6/1N5 IAV 2.5-to 15-fold (Fig. 2C), correlating well with C1q-AI activity (Supplemental Fig. 1B).

The observed VN enhancement mediated by C1q cannot be solely attributed to blocking attachment since C1q concentrations (∼5 nM) that induce negligible AI considerably enhance VN (Supplemental Fig. 1C). To test whether C1q also enhances viral-endosomal fusion inhibition (FI), we incubated C1q and anti-stem Abs at the lowest Ab concentrations causing measurable C1q-mediated hemagglutination inhibition (HI) and added human red blood cells (RBC). Once HI occurred, we washed the RBCs, measured remaining virus by NA activity and excluded samples with no detectable virus (caused by residual AI). We then triggered viral fusion by adjusting the pH to 5.0 (Fig. 2D). This revealed that C1q enhances the FI-activity of anti-stem Abs (Fig. 2E).

C1q also enhances anti-stem Ab-mediated NA inhibition against a large substrate, as determined by the soluble fetuin NA-activity inhibition (NI) assay (Fig. 2F, G).

Taken together, we have shown that C1q can enhance stem Ab VN by endowing Abs with attachment inhibition and enhancing fusion and NA inhibition activities.

### C1q does not interfere with stem Ab-mediated ADCC

It was reported that anti-HA antibodies that block viral attachment interfere with stem Ab-triggered antibody-dependent cellular cytotoxicity (ADCC) responses^29^. To study the effect of C1q on anti-stem Ab-mediated ADCC, we employed a reporter cell line (Raji) expressing luciferase controlled by the NFAT promoter, which is activated by Ab crosslinking of FcyR.

First, we tested several anti-HA-head mAbs (H36-26 Sb, IC5-4F8 Sb, Y8-1A6 Sa, H2-6A1 Sa, H2-4B1 Ca, H17-L10 Ca, H9-D3 Cb) combined with anti-stem mAbs (310-16G8, 310-16F8, Fi6) on PR8 infected MDCK Siat1.

This confirmed the strong correlation between head-specific Ab inhibition of anti-stem Ab-mediated ADCC (Supplemental Fig. 1F) and attachment inhibition (measured by hemagglutination inhibition). Remarkably, while C1q-anti-stem Ab complexes mediate AI, C1q did not inhibit ADCC activation, either in the context of target MDCK Siat1 target cells infected with PR8 or HK68 using 310-16G8, 310-16F8 or CR8020 and Fi6 mAbs (Fig. 2H). This suggests that the ability of Abs to interfere with ADCC is more subtle than is measured in HI or AI assays.

### Steric hindrance accounts for C1q-mediated enhancement of anti-stem Ab action

C1q enhancement of anti-HA Ab potency could reflect increases in steric hindrance and/or Ab-Ag binding. Adding C1q to mAbs did not increase their avidities for plate-bound virus in an ELISA (Fig. 2I), implicating steric hindrance.

To directly test for C1q-mediated steric effects, we added virus to RBCs with increasing concentrations of anti-HA mAbs, incubated for 60 min at room temperature and washed the RBCs to remove excess virus and Ab. We then added C1q and let the RBCs resettle. The washing step alone did not affect HI activity, but adding C1q enabled HI for all anti-stem mAbs tested for PR8 and HK (Fig. 2J).

These findings conclusively demonstrate that C1q binding to stem-bound Ab sterically blocks viral attachment and indicates that its functional effects on anti-HA Abs are likely due to steric effects and not enhanced Ab avidity.

**Figure 2.**
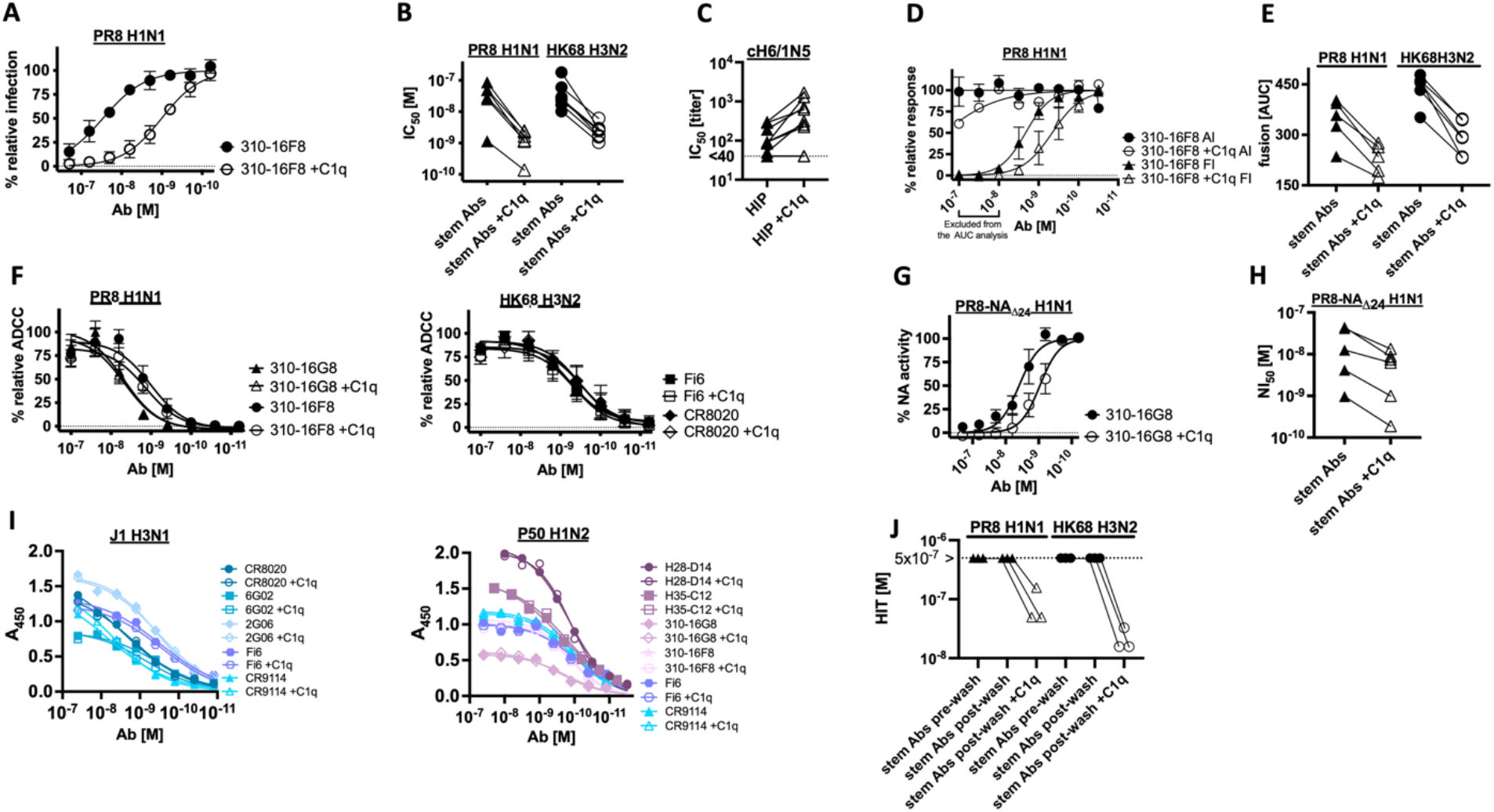
C1q enhances Anti-stem Ab VN but doesn’t interfere with ADCC. **(A)** Single-cycle flow cytometry-based VN activity *vs.* PR8 of the indicated anti-stem mAb in the absence and presence of C1q. **(B)** Effect of C1q on IC_50_ for anti-stem mAbs in single cycle VN assay using PR8 or HK. **(C)** As for B for human plasma against chimeric IAV with H6 head and H1 stem. **(D)** AI and FI activities of anti-stem Abs with and without C1q against PR8 in the attachment-hemolysis inhibition assay. **(E)** Anti-stem mAbs FI against PR8 or HK, after excluding Ab concentrations where C1q-AI was detected, with or without C1q . **(F)** Surrogate cell ADCC activity of the indicated anti-stem Ab in the absence or presence of C1q against PR8- or HK-infected cells. **(G)** NI activity of anti-stem mAb 310-IG8 with and without C1q in the soluble-fetuin neuraminidase inhibition assay **(H)** NI_50_ (Ab concentration required to inhibit the 50% of NA activity) with or without C1q for a five anti-stem Abs against the PR8-NA _Δ24_ H1N1 virus. **(I)** ELISA-determined anti-stem mAb binding with or without C1q to PR8 or HK HA expressing virions **(J)** Sequential HI for the anti-stem mAbs indicated adding C1q after washing Ab-virus bound RBCs as shown. Each experiment shows the mean of four-six technical replicates acquired in two-three independent experiments. Error bars indicate the SD. NVC and NoAbC were included in each experiment. Data are normalized using NVC set to 0% and NoAbC and/or plateau response observed for at least two highest Ab dilutions employed set to 100%. Data are plotted with GraphPad Prism7 software to generate nonlinear regression curves and calculate IC_50_, and NI_50_ using the normalized dose-response inhibition model. For the ELISA, we employed specific binding with hill slope model.

### C1q alters the BN-HA stem Ab viral escape repertoire

How does the C1q enhancement of anti-stem Ab functionality affect the HA escape repertoire? Nearly all reported anti-stem Ab escape mutants are located at or near their binding sites^30^. Does C1q extend escape residues to other locales?

As reported elsewhere for anti-stem Abs^31^, we failed to select escape mutations via single-step selection using saturating or substaturating concentrations of 310-16G8, with or without C1q. This is surprising, as single residue substitutions in the epitope encoded by point mutations are sufficient to enable escape, implying that epistatic mutations in the HA or other viral gene products are required to restore viral fitness to 310-16G8 escape mutants, as reported for other stem Abs ^32, 33^. As in prior stem mAb escape studies^32, 34, 35^ we multiply passaged PR8 with a sub-neutralizing concentration of 310-16G8, in our case with and without C1q, increasing Ab concentration with passage. As a control, we passaged PR8 without Ab with and without C1q (Fig. 3A summarizes the selection strategy). After five rounds of selection, we deep-sequenced the HA gene segment and determined the positions (H3 numbering) and frequencies of HA variants. After selection with C1q or no Ab, we failed to detect mutants at levels above the 1% sequencing error rate. Selection with Ab in the absence of C1q generated 4 non-synonymous mutants, each present at less than 2% of total sequences (HA1: N41T-1.5%, HA2: F110L-1.3%, F138Y-1.96%, D109A-1.46%) (Fig. 3B). All substitutions are present in the stem domain. N41 (Fig.3D left), present in CR6161 H1 epitope^36^, is, surprisingly, the only surface exposed escape residue. The other residues are buried in the core-facing trimer interface (D109, F110, F138) (Fig. 3D middle).

**Figure 3.**
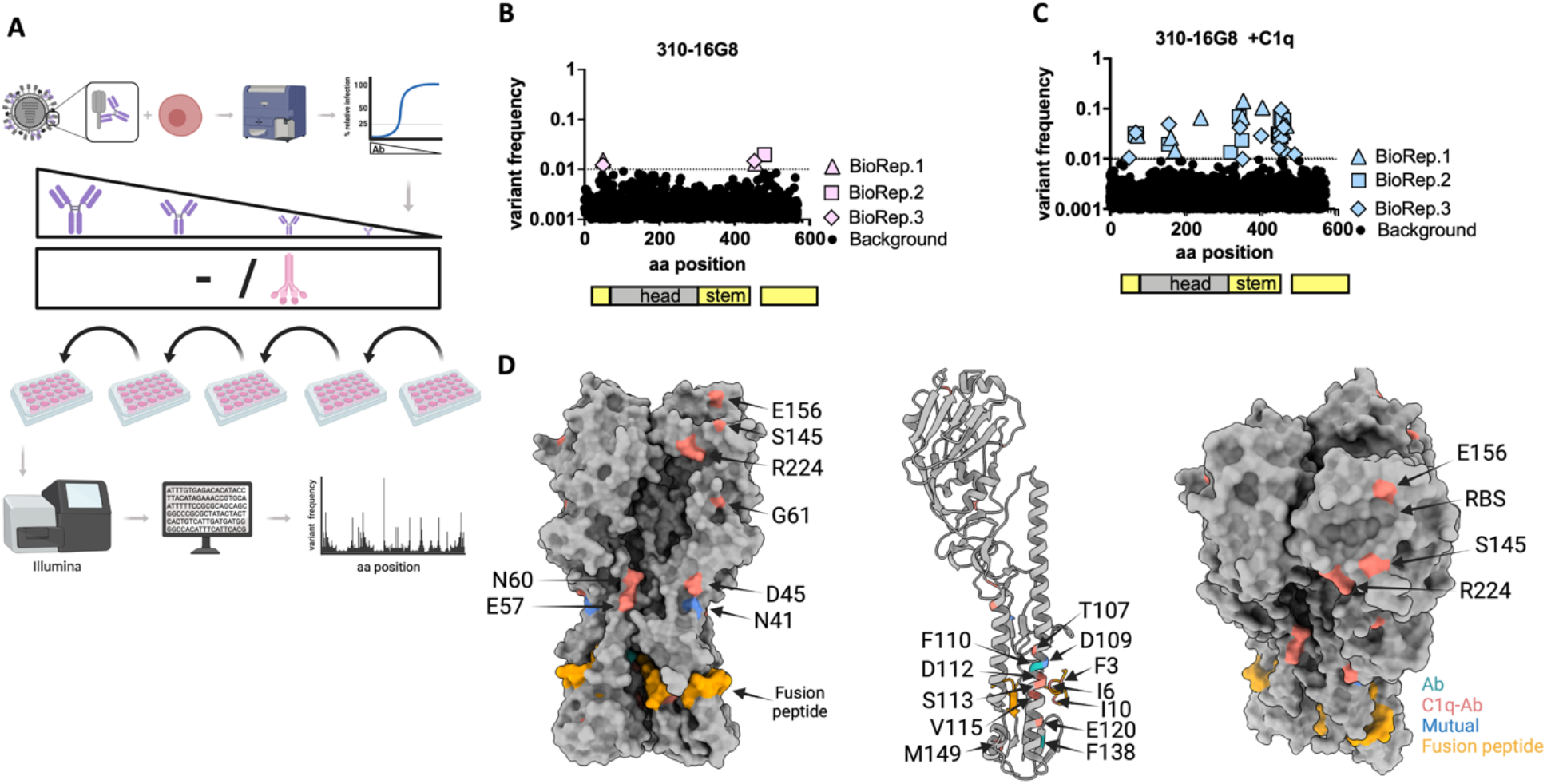
C1q-anti-stem Ab viral escape repertoire. **(A)** Selection schematic. **(B,C)** Frequencies of escape variants selected by five passages with 310-16G8 Ab in the absence (left) or presence (right) of C1q in three biological replicates. Background sequencing error rate of 1/1000 indicated by dotted line. **(D)** Identified amino acid substitutions rendered on the PR8 HA trimer structure (PDB: 1RU7) using UCSF ChimeraX_Dially. Left, trimer side view. Middle, HA monomer ribbon structure. Monomer emphasizes mutations localized at the HA stem alpha helix-C forming trimer interface. Right, tilted trimer, highlighting adsorptive mutations in or near RBS. Note that P186S was not selected by anti-stem mAb but was used as an independent test of adsorptive mutant escape.

Paradoxically, the C1q-enhanced functionality of 310-16G8 resulted in increasing both the frequency and diversity of escape mutants (Fig. 3C, Table 1), with some of the 23 detected mutants present at above 10% of the population (HA2: I6F-14.2%, E57K-10.6%). None of the Ab-only substitutions described above are present in the repertoire (though HA1 N41 is mutated to K rather than T), which extend far beyond the epitope, with several close to RBS (Fig. 4D right). The nearly identical frequencies for distant residues within the exact biological replicate (HA1-R224I-HA2-F3L, HA2-F3L-E120K, HA1-G61R-HA2-D109G, HA2-N60Y-D109G) are consistent with the selection of double mutations within a given clone (likely epistatic, to be investigated in future experiments).

**Table 1.** Summary of identified variants.

| C1q | Variant (H3 #) | Subunit | Frequency |  |  |
| --- | --- | --- | --- | --- | --- |
|  |  |  | BioRep1 | BioRep2 | BioRep3 |
| No | **N41T** | HA1 | 1.5% |  | 1.22% |
| No | **D109A** | HA2 |  |  | 1.46% |
| No | **F110L** | HA2 | 1.3% |  |  |
| No | **F138Y** | HA2 |  | 1.96% |  |
| Yes | **\*N41K** | HA1 | *2.7% |  |  |
| Yes | **⊕G61R** | HA1 | 2.96%. | ⊕3% | 3.4% |
| Yes | **\*S145N** | HA1 | *2.65% | 1.88%. | 5% |
| Yes | **E156K** | HA1 | 1.46% |  |  |
| Yes | **◇R224I** | HA1 | ◇6.67% |  |  |
| Yes | **◇⊠F3L** | HA2 | ◇6.7% | 6.8% | ⊠4.3% |
| Yes | **I6F** | HA2 | 14.2%. |  |  |
| Yes | **I10V** | HA2 |  | 2.28% |  |
| Yes | **D45G** | HA1 |  |  | 1.07% |
| Yes | **E57K** | HA2 | 10.56% |  |  |
| Yes | **⊙N60Y** | HA2 |  |  | ⊙2.98% |
| Yes | **T107P** | HA2 |  |  | 1.66% |
| Yes | **⊙⊕D109G** | HA2 |  | ⊕2.98% | ⊙3% |
| Yes | **D112N/Y** | HA2 | 3.46%/4.9% |  |  |
| Yes | **S113T/L/T** | HA2 |  | 2.8%. | 1.49%. 9.7% |
| Yes | **V115M** | HA2 |  | 6.43% | 2.54% |
| Yes | **E120K. G/ ⊠K** | HA2 | 4.63%. | 5.65%. | 1.5%/⊠4.3% |
| Yes | **M149K** | HA2 |  |  | 1.25% |
| Yes | **S163L** | HA2 |  | 1% |  |
| Yes | **V309I** | HA1 |  | 1.3% |  |

\*◇⊕⊠⊙ symbols indicate different mutations present at near identical frequencies in the same sample

Using reverse genetics, we generated 13 representative escape mutants. All mutants reached similar infectivity titers following growth in MDCK cells (Supplemental Fig. 2B). By ELISA, none of these mutants demonstrated a significant loss in affinity to the 310-16G8 selecting mAb (Supplemental Fig. 2), though HA2 D112 showed a decrease in maximal antibody binding at saturation. Using a one-step VN assay to measure entry inhibition, all mutants demonstrated escape from 310-16G8 + C1q relative to *wt* virus (Figure 4), confirming that the mutations are *bo na fide* escape mutations.

**Figure 4.**
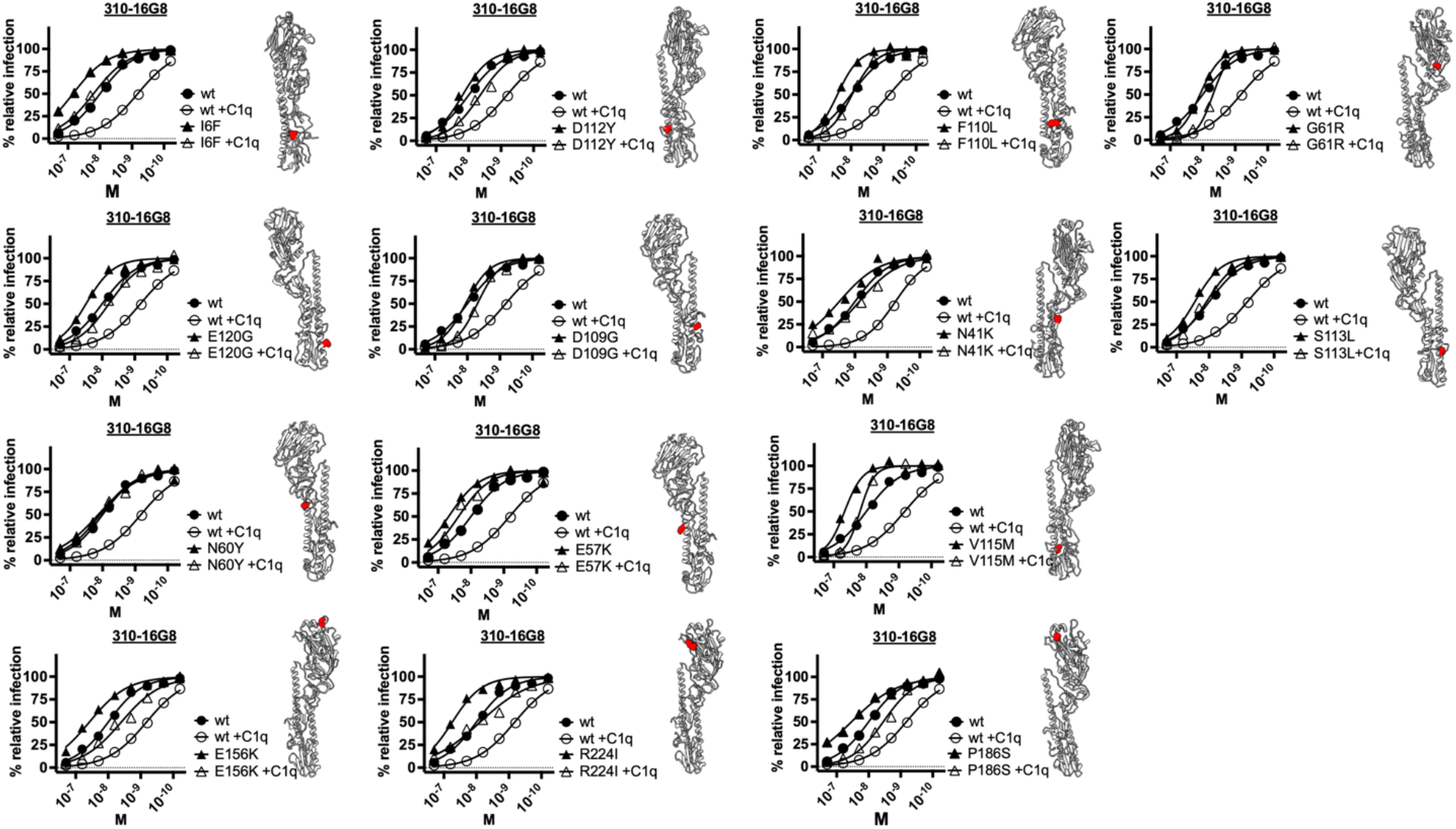
Escape mutations provide resistance to anti-stem Ab-C1q virus neutralization. We determined the VN activity of the 310-16G8 anti-stem Ab with and without C1q against the indicated PR8 virus escape variants. Locations of amino acid substitutions on HA monomers are shown in red. Each experiment shows the mean of four-six technical replicates acquired in two-three independent experiments. Error bars indicate the SD. NVC and NoAbC were included. Data are normalized using NVC set to 0% and NoAbC and/or plateau response observed for at least two highest Ab dilutions employed set to 100%. Nonlinear regression curves were generated with Prism software using the normalized dose-response inhibition model.

Notably, some residues present in C1q Ab escape mutants (HA1 E156K, R224I) are known to increase HA binding to SA receptors^49^ (though R224 is likely to be an epistatic mutation improving HA2 F3L fitness). We confirmed that increased HA receptor avidity contributes to virus escape from C1q-stem Ab VN by including the classic PR8 HA1 P186S adsorptive mutant in the analysis (lower right-hand panel), whose increased receptor binding enables escape from polyclonal Ab *in vitro*^50^ and *in vivo*^49^ despite being nearly antigenically identical to *wt* virus^50^.

These findings demonstrate that C1q expands the anti-stem Ab escape repertoire to include residues located throughout HA, including residues that escape VN by increasing HA receptor affinity.

### Virion antigen geometry modulates C1q-Ab activity

PR8 virions possess ∼ 400 HA trimers and 40 NA tetramers, present in clusters in close proximity to HA as revealed by EM images of virions^37^ and anti-HA Ab inhibition of NA activity^10, 38^. Does C1q enable anti-NA mAbs to block attachment? We tested a panel of NA mAbs at constant concentration (200nM) with graded amounts of C1q (starting at 220nM) in the muNANA-based AI assay (anti-NA Abs do not inhibit cleavage of small substrates). None of four NA-specific mAbs exhibit C1q-AI (Fig. 5A), consistent with the idea that virion surface protein geometry is a critical factor affecting C1q steric effects.

**Figure 5.**
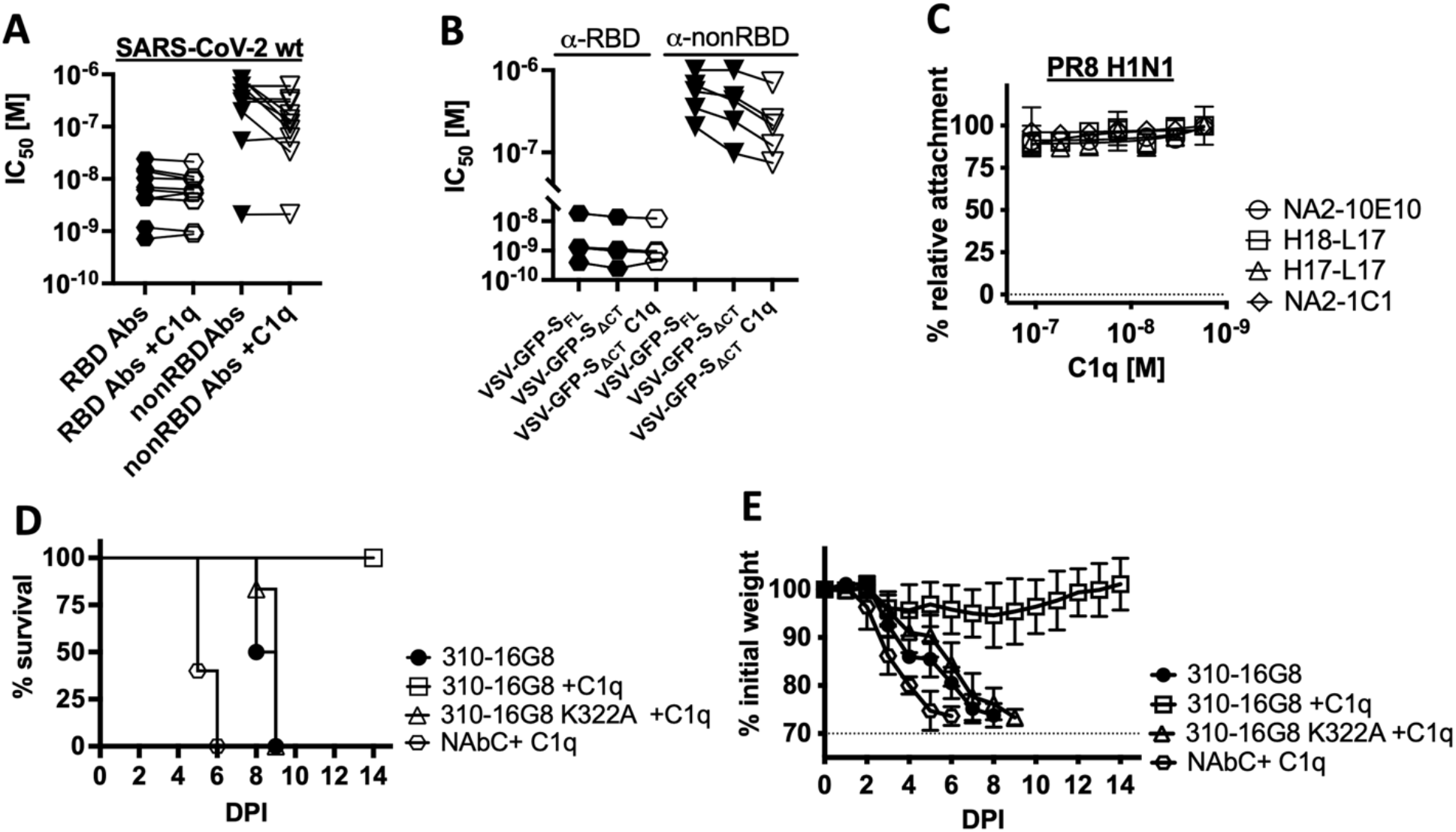
Virion antigen geometry modulates C1q-Ab activity. **(A)** AI activity of NA-specific mouse mAbs in the presence of C1q in the fetuin-attachment assay. We employed a constant Ab concentration (200nM) and diluted C1q (starting at 110nM). **(B)** IC_50_ for 11 SARS-CoV-2 Spike RBD and 11 non-RBD specific mAbs against self-reporting (GFP expressing) SARS-Cov-2 virus in a one-step flow cytometric VN assay. **(C)** IC_50_ for five Spike RBD and five nonRBD specific mAbs against VSV-GFP-S_FL_ or VSV-GFP-S_ΔCT_ GFP reporter pseudoviruses in one-step flow cytometric VN assay. Alternatively C1q was combined with mAbs. **(D)** The initial body weight change after one dose intranasal delivery of Abs indicated with or without C1q and lethal IAV challange **(E)** Fraction survival after one dose intranasal delivery of Abs indicated with or without C1q and lethal IAV challange

We extended these findings to SARS-CoV-2 Spike protein by testing a panel of 22 published mAbs (11 anti-RBD, 11 anti-non-RBD, Supplemental Tab. 2)^39–41^ for SARS-CoV-2 VN activity in the presence and the absence of C1q. While anti-RBD Abs exhibited negligible C1q-VN changes, C1q enhanced the VN activity of five non-RBD Abs three-to seven-fold (Fig. 5B). Notably, all the enhanced Abs belong to the same competition cluster^40^, indicating that this antigenic site supports an Ab orientation that enables C1q enhancement.

Why do all HA-specific mAbs but only some Spike-specific Abs enhance VN? To test whether this is related to the average distance between HA *vs.* Spike on the virion surface (11nm^42^ *vs.* 24 nm^43, 44^) we generated pseudotyped VSV recombinants with different SARS-CoV2 Spike densities using *wt* Spike and Spike_ΔCT_, which lacks the Spike C-terminal ERGIC retention sequence^45^.

Immunoblotting sucrose purified VSV confirmed that the *wt* Spike is incorporated at half Spike_ΔCT_ levels (Supplemental Fig. 1G). Testing five RBD and five non-RBD Abs (those enhanced by C1q in Fig. 5B) for VN activity against the two viruses (Fig. 5C) revealed that C1q does not enhance RBD-specific mAbs regardless of Spike density. By contrast, for the five non-RBD-Abs, doubling spike density alone enhanced VN, the phenomenon further boosted with C1q.

These findings indicate that Spike density can influence Ab VN activity, including the potentiating effect of C1q, and support the importance of using SARS-Cov2 or close proxies for VN assays.

### C1q enhances anti-stem Ab-mediated protection against lethal mouse IAV infection

To extend our findings to an *in vivo* model, we administered the human anti-stem mAb 310-16G8 (300ng per mouse) intranasally to B6 mice with or without hC1q (25µg per mouse). As a control, we including the C1q binding-deficient Fc-K322A Ab variant in the experiment. One hour post-treatment, we challenged mice with a lethal IAV infection dose. We monitored morbidity (body weight change, hunched posture, scruffiness, mobility), and sacrificed mice according to our challenge protocol when weight loss reached 30% of body mass (Fig. 5D, E). C1q alone did not affect IAV-associated morbidity and mortality. While Ab-alone temporarily postponed morbidity and prolonged survival (72 H), 100% of animals succumbed to the infection. This is most probably due to the incompatibility and/or local concentration of mouse C1q, an earlier one supported by the inability of hIgG1 to bind recombinant mouse C1q as determined by BLI (Supplementary Fig. 2C) Importantly, in the presence of human C1q, the non-protective anti-stem Ab dose conferred 100% protection with mild body weight loss (max 6%) and minor signs of illness. Moreover, K322A Ab closely resembled wt Ab-alone treatment even if C1q was co-administered, conclusively demonstrating the relevance of our observations for C1a enhanced protection of anti-stem Abs against respiratory IAV infection.

## Discussion

C1q in blood and body fluids binds avidly to IgG antibodies present in a geometric array on viruses or other multivalent antigens to enable multivalent binding. Here, we confirm findings from the Gerhard lab that C1q enhances the ability of Abs specific for the HA globular domain to block viral attachment and infectivity^23^, with the effect increasing with Ab epitope distance from the HA RBS. We extend these findings by demonstrating the remarkable ability of C1q to enable anti-stem Abs to block virus attachment and enhance inhibition of HA-mediated fusion and NA receptor cleavage.

We provide evidence supporting the *in vivo* relevance of C1q enhancement of stem Ab VN activity using the mouse influenza model. We chose intranasal administration of human mAb with human C1q to mimic human respiratory mucosal immunity. Relative to prior studies^46, 47^ (which used 300-100µg mAb per mouse, given parenterally), much less mAb (300ng per mouse) was required to achieve protection using intranasal Ab induction, likely due to improved mucosal vs. systemic immunity ^48^. Reported levels of C1q in healthy volunteer human bronchiolar alveolar fluids range from very low ^49^ to 29% of serum levels^50^, the latter consistent with C1q function in mucosal immunity to initial infection. Even when C1q is present in non-physiological levels initially, this will almost certainly change with inflammatory increases in vascular permeability induced by viral infection.

It is counter-intuitive that in enhancing anti-stem Ab function (even providing novel functionality in conferring AI), C1q substantially expands the escape repertoire to include residues distant from the selecting Ab epitope. We directly show that many of these substitutions (Supplementary Fig. 2A) do not reduce selecting Ab avidity while still providing escape from stem Ab-mediated entry VN, pointing to alternative escape mechanisms. Many substitutions are present in the stem, but distant from defined stem epitopes (Fig. 3D)^47, 51^. T107, D109, D112, S113, V115, E120, and M149 are located in the HA stalk core helixes C-D^30^, and might modulate HA flexibility and/or fusion pH. Indeed, three substitutions, HA2 F3L, HA2 I6F, HA2 I10V, are in the fusion peptide itself and are known to affect fusion^52, 53^. We previously showed that substituting PR8 HA2 F3 alters fusion and HA flexibility, as demonstrated by VN resistance to a Sa site-specific mAb (Y8-10C2) whose epitope is only exposed in trimers at elevated temperature^48^. This suggests that changes in HA breathing contribute to escape from C1q anti-stem Ab VN, as will be explored in future experiments.

Several substitutions are known to increase receptor avidity (E156K, R224I)^54, 55^, consistent with viral attachment-based selection. We show that the classic P186S adsorptive mutant^50^ also demonstrates escape from C1q enhanced-anti-stem Ab VN. Surprisingly, each of the three adsorptive mutants escapes VN without C1q in a one-step VN assay, providing the first evidence that receptor avidity modulates what appears to be strictly fusion-based Ab-mediated entry inhibition. Why such mutations are not selected without C1q is an interesting question for future studies.

Lee *et al.* ^33^ recently reported that adsorptive mutations restore fitness to anti-stem mAb escape mutations localized in the selecting Ab epitope. Presumably, epistasis, in this case, is due to alterations in HA receptor binding due to long-range conformational effects. We note that these findings are related to our own, but only indirectly, since we document the direct effects of adsorptive mutations in escaping stem Ab VN.

Regardless of the exact mechanism by which substitutions enable stem Ab+C1q selection, these findings reinforce our previous studies^6, 54, 56, 57^ showing that inferring selective pressures that lead to the emergence of given escape mutations is difficult, if not impossible, particularly in real-world virology. This is underscored by the C1q-stem mAb selection of head mutations (E156K, G61R, S145N) known to affect HA antigenicity^3^.

Extending our findings to SARS-CoV2, we found that none of the anti-RBD or anti-NTD Abs tested exhibit significant C1q-VN enhancement. As with HA tip-specific Abs (Sa, Sb), anti-RBD Abs probably exhibit close to maximal steric hindrance in blocking attachment, limiting C1q contribution. We show that VN mediated by five non-RBD specific mAbs is enhanced by C1q, extending the phenomenon to another important human virus and that the low Spike density on corona virions is likely to limit C1q function.

In summary, we have shown that C1q expands the function of classically “non-neutralizing” Abs for IAV and SARS-CoV2 and is likely to have similar effects for other viruses. Most generally, our findings underscore the importance of studying immune effectors in of the context of their evolutionary selection.

## Materials and Methods

### Cell lines and viruses

We cultured MDCK SIAT7e^58^ suspension cells on a rotatory shaker at 225RPM in DMEM (Gibco) supplemented with 8% FBS (HyClone) and 500 µg/ml gentamicin (Quality Biologicals) and 500 µg/ml geneticin (Gibco) at 37°C and 5% CO2. We cultured MDCK SIAT1 cells in DMEM (Gibco) supplemented with 8% FBS (HyClone) and 500 µg/ml gentamicin (Quality Biologicals), and 500 µg/ml geneticin (Gibco) at 37°C and 5% CO2. We cultured human embryonal kidney 293T cells in DMEM supplemented with 8% FBS (HyClone) and 500 µg/ml gentamicin at 37°C and 5% CO2. We used Expi293 cells (Thermo Fisher) for monoclonal Ab expression. We propagated and cultured the cells as recommended by the manufacturer. We cultured BHK21 and BHK21-Ace2^59^ cells in DMEM (Gibco) supplemented with 8% FBS (HyClone) and 500 µg/ml gentamicin (Quality Biologicals), and 500 µg/ml geneticin (Gibco) at 37°C and 5% CO2. In the case of BHK21-Ace2, in every fourth-fifth passage, we added hygromycin gold at 250µg/ml (Invitrogen).

We employed the following viruses in this study: A/Puerto Rico/8/34/H1N1 (Walter Gerhart), A/Hong Kong/1/68 (H3N2), P50 (H1N2 A/Puerto Rico/8/34/H1 x A/Hong Kong/1/68/N2), J1 (H3N1 A/Hong Kong/1/68/H3 x A/Puerto Rico/8/34/N1), chimeric HA cH6/1N5 x A/Puerto Rico/8/34-core, self-reporting replicative A/Puerto Rico/8/34/H1N1x NS1mCherry^10^ A/Puerto Rico/8/34/H1N1/NA_Δ24_ (PR8-NA_Δ24_)^10^. We generated a reassorted self-reporting A/Hong Kong/1/68 (H3N2) x A/Puerto Rico/8/34/H1N1xNS1mCherry by cotransfection of the six core influenza gene–coding plasmids (including pDZ-NSmCherry) and pDZ-HA (HK68) and pDZ-NA (HK68) as published previously^10^. To generate HA variants, we performed site-directed mutagenic PCR with 5’ phosphorylated primers listed in Supplemental table 2. We used the PR8 pDZ-HA plasmid as a template and Kod Hot start DNA polymerase (Millipore Sigma). We followed the manufacturer’s recommendations to PCR mutagenize the pDZ-HA open reading frame. We recircularized linearized plasmid PCR products with a Rapid DNA Ligation Kit (Roche) and transformed plasmids into DH5α-competent cells (Thermo Fisher). We sequenced plasmids and confirmed mutations after purification with plasmid purification (Qiagen). For virus purification, we clarified allantoic fluid at 4,700 rounds per minute (RPM) for 10 min and pelleted the virus by centrifugation for two h at 27,000 RPM. We incubated pellets overnight in 2 ml PBS with calcium and magnesium (PBS++). We purified the virus by centrifugation on a discontinuous 15–60% sucrose gradient, collecting the virus at the interface, and pelleting 34,000 RPM for two h. After resuspending the pellet in 500 µl PBS++ overnight, we measured total viral protein with the DC Protein Assay (Bio-Rad). The following reagent was obtained through BEI Resources, NIAID, NIH: SARS-Related Coronavirus 2, Isolate USA-WA1/2020, Recombinant Infectious Clone with Enhanced Green Fluorescent Protein (icSARS-CoV-2-eGFP), NR-5400. The virus was propagated and characterized by the National Institute of Allergy and Infectious DiseasesSARS-CoV-2 Virology Core.

We generated non-replicative self-reporting VSVΔG-EGFP-SARS2-S_FL_ or Sι1_CT_ as described previously^59^. The following reagent was produced under HHSN272201400008C and obtained through BEI Resources, NIAID, NIH: Vector pCAGGS Containing the SARS-Related Coronavirus 2, Wuhan-Hu-1 Spike Glycoprotein Gene, NR-52310. The open reading frame of Spike glycoprotein (Sgp), excluding the last 16 amino acids of the carboxy terminus, was amplified in PCR using Q5® Hot Start High-Fidelity 2X Master Mix (New England Biolabs) using primers flanked with *SfiI* restriction enzyme recognition sequence (underlined) followed by irrelevant nucleotide to facilitate the digestion of PCR product. Primer sequences are present in table 2. The PCR product was purified using DNA Clean & Concentrator-5 (Zymo Research), sequentially digested with FastDigest *SfiI* and FastDigest *DpnI* (Thermo Scientific), and the cloning insert purified by DNA Clean & Concentrator-5 (Zymo Research). Vector pSBbi-bla, a kind gift from Eric Kowarz (Addgene plasmid #60526), was chosen to generate SgpΔ16Cter expressing plasmid. FastDigest SfiI digested plasmid pSBbi-bla (Thermo Fisher Scientific), dephosphorylated using FastAP Thermosensitive Alkaline Phosphatase (1 U/µL) (Thermo Scientific), size-selected by 1% agarose electrophoresis, and purified using Zymoclean Gel DNA Recovery Kit (Zymo Research). Prepared insert and vector fragments were ligated using Rapid DNA Ligation Kit (Roche), transformed to One Shot TOP10 Chemically Competent *E. coli* (Invitrogen), and selected on carbenicillin LB plates. The sequence of picked clones was confirmed by Sanger sequencing. We employed a commercial VSVΔG-EGFP system (Kerafast). Briefly, we seeded a T75 tissue flask to 60-70% confluency (five to seven million 293T cells) in DMEM(Gibco) supplemented with 8% FBS (Hyclone) and 10mM HEPES (Corning). We combined 50 μL FuGene low cytotoxicity transfection reagent (Promega) and 2 mL OPTIMEM media (Gibco). We combined 32 μg of pCAGGS-S-Covid19 (S_FL_) (BEI NR-52310) or pSARS2-SΔ_CT_ and 4 μg pTRMPSS2 (VRC/NIAID) diluted in 2 mL OPTIMEM media (Gibco) (DNA OPTIMEM mixture). After five minutes of incubation at RT, we incubated the FuGene and pDNA OPTIMEM mixtures at room temperature for 10-15 minutes. Media from the seeded T75 flask was gently aspirated and replaced with 20 ml of the fresh DMEM (Gibco) supplemented with 8% FBS (Hyclone) and ten mM HEPES (Corning). The transfection mixture was added dropwise and incubated with the cell culture for 24-48 hours at 37°C in a CO2 incubator. The virus mixture was prepared by combining 0.3 mL of VSVΔG-EGFP-Gp P3 (titer ∼10e8/mL) and 1.7 mL of DMEM (Gibco) supplemented with 8% FBS (Hyclone) and ten mM HEPES (Corning). The medium was aspirated from the transfected T75 flask cell culture, and the virus was added and incubated at 37°C for one hour in a CO2 incubator while gently rocking every 10-15 minutes. Following incubation, we aspirated media, and we added 20 mL of DMEM (Gibco) supplemented with 8% FBS (Hyclone) and ten mM HEPES (Corning). We inspected cells by fluorescence microscopy, and once most of the cells turned green-fluorescent and rounded (30-36 hours), we harvested media. We centrifuged the suspension at 4,700 RPM for 10 minutes at four °C and transferred the supernatant. We resuspended pelleted cells in 1-2 ml PBS, freeze-thawed three times, and sonicated on the ice at 50% amplitude five times in one-second pulses. We centrifuged the mixture at 4,700 RPM for 10 minutes at 4d°C. We collected the supernatant, combined it with media supernatant, and snap-frozen it in an isopropanol-dry ice mixture for 20 minutes before storing it at -80°C. We titrated VSVΔG-EGFP -SARS2-Sgp (S_FL_ and Δ_CT_) on BHK21-ACE2 cells by flow cytometry. To exclude the contribution of potentially recycled VSV-Gp, we compared the susceptibility to infection of wtBHK21 and BHK21-ACE2 cells and infectious titer in the presence and absence of the 1E9F11 and 8G5F11 anti-VSV-Gp antibody mixture (both 10 μg/mL). We concluded that the effect of potentially recycled VSVGg is negligible for the VSVΔG-EGFP-SARS2-Sgp (S_FL_ and Δ_CT)_ virus stocks.

### Escape variants selection

For the 310-16G8 Ab, we seeded 80,000 MDCK SIAT1 cells per well in 24 well cell culture-treated plates (Corning). We combined roughly 10^3^ TCID_50_ of PR8 per well in 150 μl of infection media supplemented with 310-16G8 (2.7 nM) or 310-16G8 and C1q (160 nM). After one hour of incubation at 37°C, we washed cells twice with DPBS and added Ab-virus or C1q-Ab-virus mixture for two hours at 37°C. Alternatively, we incubated the virus in infection media or supplemented it with C1q. We washed the cells three times with DPBS to remove the uninternalized virus, and we added 300 μl of infection media supplemented with 310-16G8 (2.7 nM) or 310-16G8 and C1q (160 nM). We inspected the plate three to four days post-infection and collected 150 μl from infected wells. We added 310-16G8 (4.6 nM) or 310-16G8 and C1q (160nM). After one hour of incubation at 37°C, we transferred 150ul of media to fresh cells for two hours. As mentioned above, we washed and replaced infection media with Ab, Ab, and C1q. We repeated passaging for three more cycles while increasing the 310-16G8 Ab concentration (P3-6.6 nM, P4-10 nM, P5-13 nM). We used supernatants collected from selection experiments for sequencing of viral populations.

### Virus sequencing

We extracted influenza virus genomic RNA from the clarified allantoic fluid using the QIAamp Viral RNA Mini Kit (Qiagen) according to the manufacturer’s protocol. We amplified complete genomes with a multiplex RT-PCR protocol using SuperScript III One-Step RT-PCR System with Platinum Taq DNA Polymerase and primers targeting conserved five ʹ and 3ʹ sequences in the viral RNA^60^. Libraries for next-generation sequencing were constructed from 1-ng amplified genomes using the Nextera XT DNA Library Preparation Kit (Illumina) and sequenced using the MiSeq Reagent Kit v3 on the MiSeq Platform (Illumina). Viral sequences were assembled de novo using a high-throughput assembly pipeline^61^, and variant statistics were assembled using custom Perl scripts.

### Antibodies

We produced the human IgG1 mAbs 310-16G8, 310-18F8, 310-18E7, 310-18D5, CR6261, FI6, CR9114, SFV005 2G02, 008-10053 6C05, 042-2F04, 017-100116 5B03, and CR8020^35, 62–64^ by transient plasmid cotransfection in the Expi293 Expression System (Thermo Fisher) following the manufacturer’s recommendations. Following the manufacturer’s recommendations, we purified Abs using HiTrap Protein G HP antibody purification columns (Cytvia) and Akta Start protein purification system (Cytvia)Chimeric. Abs’ H28-D14 and H35-C12 expression plasmids were custom-made (VectorBuilder) after we determined sequences of VH and VL portions from in-house mouse hybridomas. To generate germline reverted H28-D14 and 310-16G8 Abs’ avidity variant, we performed site-directed mutagenic PCR with 5’ phosphorylated primers listed in Supplemental table 2. We used the pH28D14 HC and LC plasmids as templates and Kod Hot start DNA polymerase (Millipore Sigma). We followed the manufacturer’s recommendations to PCR mutagenize the VH and VL gene segments. We re-circularized linearized plasmid PCR products with a Rapid DNA Ligation Kit (Roche) and transformed plasmids into DH5α-competent cells (Thermo Fisher). We sequenced plasmids and confirmed mutation after purification with plasmid purification (Qiagen). We expressed and purified Abs as described above. SARS2 Spike human IgG1 monoclonal RBD Abs COVA1-18, COVA2-39, COV2-2832, COV2-2807, COV2-2780, C102, C119, C105, C110, C121, C135, and nonRBD Abs COVA1-25, COVA1-02, COVA1-06, COVA1-09, COVA2-10, COVA2-30, COVA1-12, COVA2-28, COVA2-33, COVA2-42, COVA1-22 (NTD)^39–41^, were custom produced based on available VH, VL sequences (Genescript, Echo Biosolution, BioinTron). The PR8 HA mAbs H2-6A1 (Sa), Y8-1A6 (Sa), H36-4 (Sb), H28-E23 (Sb), H2-4B1 (Ca), H17-L10 (Ca), H17-L7 (Cb), HA1-specific mAb CM1, PR8 NA Abs NA2-10E10-mIgG2a, H18-L17-mIgG2b, H17-L17-mIgG2a, NA2-1C1-IgG1, NP-specific mAb HB65 conjugated with AF 647, were prepared in the laboratory and/or published previously^65^. We used rabbit polyclonal anti-PR8 HA antibody (Sino Biological), SARS-CoV-2 Spike S2 Subunit Antibody (R&D systems), IRDye® 800CW Goat anti-Human IgG (H + L), IRDye® 800CW Goat anti-Mouse IgG (H + L), IRDye® 800CW Goat anti-Rabbit IgG (H + L), Goat F(ab)2 anti-human IgG: HRP (Biorad), Goat anti-mouse IgG (H/L): HRP (Biorad)

### Virus hemagglutination and hemagglutination inhibition assay

We diluted allantoic fluid containing half-log serial dilutions of virus in Dulbecco’s PBS (DPBS) in a round-bottom 96-well plate (Greiner Bio-One). We combined 50 µl of the virus dilutions with 50 µl 0.5% turkey RBCs and incubated at four °C for one hour. We determined HA titer as the reciprocal of the highest dilution providing full hemagglutination. We tested all viruses in triplicate in two independent experiments.

We serially diluted Abs in DPBS in the round bottom 96-well plate (Corning) for the HI. We combined 25 µl Ab dilutions with 25 µl IAV (4HAU) diluted in DPBS, and we incubated the mixture at 37°C for one hour. We combined 50 µl of the Ab-virus dilutions with 50 µl 0.5% turkey RBCs and incubated at room temperature for one hour.

Alternatively, for the sequential HI, we serially diluted Abs (twofold starting at 500nM) in 10 µM oseltamivir impurity B CRS (European Pharmacopoeia Reference Standard) in DPBS or ten µM oseltamivir impurity C, C1q (160 nM) in DPBS. We combined 25 µl Ab dilutions with 25 µl IAV (4HAU) diluted in 10 µM oseltamivir impurity B CRS in DPBS, and we incubated the mixture at 37°C for one h. We combined 50 µl of the Ab-virus or C1q-Ab-virus (control) dilutions with 50 µl 0.5% turkey RBCs and incubated at room temperature for one hour. We scanned the plates. For the second round of HI, we washed immunocomplexes three times by adding 150 µl 10 µM oseltamivir impurity B CRS in DPBS, gently pipetting mixtures ten times, and spinning plates at 2,000RPM for five minutes. We resuspended immunocomplexes in 100 µl of 10 µM oseltamivir impurity B CRS in DPBS or ten µM oseltamivir impurity B CRS, C1q (160 nM) in DPBS, and we incubated the mixture at 37°C for one h. We rescanned the plates.

### In-cell immunoassay and median tissue culture infectious dose (TCID_50_)

We performed assay as described previously^10^. We seeded MDCK SIAT1 cells (10,000–50,000 per well) in a 96-well plate (Costar). The following day we washed cells with DPBS twice and added half-log10 dilutions of the virus sample in infection media (MEM medium (Gibco) containing 0.3% BSA (fraction V; Roche), 10 mM HEPES (Corning), 500 µg/ml gentamicin (Quality Biologicals), and 1 µg/ml TPCK trypsin (Worthington)). At 18 h p.i., we fix-permeabilized the cells with 100% ice-cold methanol for 20 min. We washed the cells with DPBS and added 30 µl HB65 anti-NP mAb diluted in Odyssey Blocking Buffer PBS (OBB; Li-Cor) for 1 h at room temperature. We washed the plate with DPBS containing 0.05% NP-40 (Thermo Fisher) twice and added IRDye 800CW goat anti-mouse Ab (Li-Cor) 1,000× diluted in OBB. After 1-h incubation at room temperature, we washed the plate twice as described above with one final milliQ water washing step. We scanned the plates using a near-infrared imaging system (OdysseyCLx; Li-Cor). We used ImageStudioLite software to quantify the measured signal. For the positive/negative threshold, we used doubled averaged signal of eight uninfected wells. We used the Spearman and Karber method to calculate TCID_50_. We plotted TCID_50_ data using GraphPad Prism7.

### ELISA

We coated ELISA plates (half-area 96-well; Greiner Bio-One) with ten ng of respective purified whole IAV diluted in DPBS (30 µl per well). After overnight incubation at four °C, we washed plates three times with DPBS supplemented with 0.05% Tween-20. Alternatively for the HA variants, we combined 100µg of the sucrose cushion purified viruses in 250ul DPBS with 250ul of the 2% NP-40 (Thermofisher), we incubated mixture at 37°C for 30 minutes and we added 4.5ml DPBS (100x solution at final concentration 20ug.ml^-1^). To further account for coating differences, we normalized by signal acquired during thd experiment on the same plate with rabbit polyclonal anti-PR8-HA antibody (Proscience). We blocked the wells with 1% BSA (Fraction V, Roche) in DPBS (70 µl per well) for one hour at room temperature. We serially diluted (half-log_10_) Abs in 1% BSA in DPBS or 1% BSA, C1q (160nM) in DPBS and added 30 µl to plates for 90 min at 37°C. After extensive washing with DPBS+0.05% Tween-20, we detected bound Abs by incubating plates with 30 µl HRP-conjugated secondary Abs following a manufacturer’s recommendation for one h at room temperature. We washed ELISA plates with DPBS+0.05% Tween-20 and incubated them with SureBlue TMB Microwell Peroxidase Substrate (KPL) for 5 min at room temperature. We stopped the enzymatic reaction by adding 1N HCl. We measured absorbance at 450 nm (A_450_) on a Synergy H1 plate reader (Biotek) and calculated the dissociation constant (Kd) from dilution curves using GraphPad Prism 7 software to fit one-site binding.

### Fetuin attachment inhibition (AI) assay

We performed an AI assay as described previously^25^. Briefly, we coated black, half-area, high-binding 96-well plates (Nunc) with 30 µl fetuin (Sigma Aldrich) in DPBS (50 µg/ml) overnight at 4°C. We blocked the wells with 1% BSA in DPBS (70 µl per well) for one hour at room temperature. We determined the viruses’ dilutions that yield a sufficient signal (roughly 20-40HAU). We serially diluted Abs in 1% BSA in DPBS (Gibco) or 1%BSA, C1q (160 nM, 70 µg/ml) by half-log_10_ starting at 200 nM. Alternatively, we diluted C1q in 1%BSA, Ab (100/200 nM) by twofold, starting at 200nM. We combined Abs or C1q dilutions with IAV and incubated them for 60 min at 37°C. We then transferred 50 µl Ab-virus or C1q-Ab-virus mixture to washed fetuin-coated plates, incubated for one hour at four °C, and removed the free virus by washing extensively with DPBS. We added 50 µl of 200 µM muNANA in 33 mM MES, pH 6.5, with four mM CaCl_2_ and incubated for one h at 37°C. We measured fluorescence (excitation [Ex] = 360 nm; emission [Em] = 450 nm) using a Synergy H1 plate reader (Biotek).

### The free fetuin neuraminidase inhibition assay

To exclude AI contributing to NI in commonly used ELLA^25^, we modified assay to use free fetuin. We biotinylated fetuin by combining 1.5ml fetuin (Sigma Aldrich) at 25 mg.ml^-1^ with 127 µl of 100nM Biotin NHS-LC (ThermoFisher) (1:20 molar ratio). We incubated mixture ON at 4 °C while agitating 150 RPM. We removed free biotin with zeba desalting column (ThermoFisher). We combined 25 µl of PR8-NA_Δ24_ (40HAU) with 25 µl of the respective Ab with or without C1q at 140 µg.ml^-1^ and we incubated mixture for 90 min at 37 °C. We added 50 µl of biotinylated fetuin at 25 µg.ml^-1^ and we incubated mixture ON at 37 °C. We transferred 90 µl to streptavidin coated (50 µl three µg.ml^-1^) and 1 % BSA blocked 96 well plates and incubated at RT for one hour.

After washing three times with DPBS+0.05% Tween-20, we added 50 µl of peanut lectin-HRP (Sigma Aldrich) (diluted 1,000 times) at RT for one hour. We washed as above and we added 50 µl SureBlue TMB Microwell Peroxidase Substrate (KPL) for 10 minutes at RT and we stopped reaction by adding 50 µl 1N HCL. We measured absorbance at 450 nm (A_450_) on a Synergy H1 plate reader (Biotek). We plotted the data using GraphPad Prism 7 software.

### AI-linked hemolysis inhibition assay

We first titrated the Abs to yield HI on 1% human RBC in C1q (220 nM) presence against 20 HAUs of IAVs. We diluted anti-stem Abs (starting at the lowest concentration, which yielded measurable HI) in C1q-DPBS (Gibco) (220 nM). We combined 25 µl of the dilutions with 25 µl (20 HAUs) of the PR8 (H1N1) or HK68 (H3N2) virus in a round bottom 96-well plate (Corning), respectively. After one hour of incubation at 37°C, we combined 50 µl of C1q-Ab-virus mix with 50 µl of 1% human RBCs. Once hemagglutination occurred, we spin-rinsed (2000 rpm for five minutes) hRBCs thrice with 250 µl DPBS. We aspirated residual DPBS and added 40 µl of 100 µM muNANA in DPBS (since 33 mM MES, pH 6.5, with four mM CaCl_2_ caused hemolysis). After one hour of incubation at 37 °C, we measured fluorescence (excitation [Ex] = 360 nm; emission [Em] = 450 nm) using a Synergy H1 plate reader (Biotek). We spin-rinsed (2000 rpm for five minutes) hRBCs thrice with 250 µl DPBS. We aspirated residual DPBS, added 70 µl of citrate buffer (pH=5), and incubated plates for one hour at 37 °C. We spun down the plates (2000 rpm for five minutes) and transferred 40 µl of the solution to the flat bottom 96-well plates. We measured absorbance at 410 nm using a Synergy H1 plate reader (Biotek). We plotted the data using GraphPad Prism7 software.

### Human plasma samples and red blood cells

Human blood samples were obtained from healthy donors on an IRB-approved NIH protocol (99-CC-0168). Research blood donors provided written informed consent, and blood samples were de-identified prior to distribution. Clinical Trials Number: NCT00001846. We separated plasma from the blood cells by centrifugation (4,000 rpm for one minute) through the separation tubes pre-filled with separation gel (Sarstedt Serum GelZ1.1). We heat-inactivated the plasma samples at 57°C for 30 minutes, and we stored the plasma samples at -30°C.

To prepare stock (10 %) of human RBCs, we rinse and spin-washed the blood with ten volumes of the ice-cold Alsever’s solution. We aspirated residual solution, diluted RBCs to 10% in Alsever’s solution, and stored at 4°C. We freshly rinse spin-washed RBC before use and dilute it to the required concentration in DPBS.

### Antibody-dependent cell-mediated cytotoxicity

We seeded 10,000 MDCK SIAT1 cells to 96-well white tissue culture–treated plates (Costar). The following day, we washed cells twice with DPBS, and infected cells with the IAV virus (MOI = 4) diluted in the MEM media supplemented with 0.3% BSA and 10mM HEPES. At six to eight hours p.i., we washed the cells twice with DPBS and added 25 µl of the mixture of anti-stem and HA head Ab (each 200 nM) diluted in RPMI 1640 medium supplemented with 0.3% BSA. After 30 minutes of incubation at 37°C in 5% CO_2_ we added 50,000 ADCC-RL cells (Jurkat cell line expressing luciferase gene under the control of the NFAT response element and stably expressing human FcγRIIIa V158; Promega) in 50 µl RPMI 1640 medium with 6% low-IgG serum (Promega). After overnight incubation at 37°C in 5% CO2, we added 50 µl of Bright-Glo Luciferase Assay lysis/substrate buffer (Promega) and measured luminescence within 10 min using a Synergy H1 plate reader (Biotek). Alternatively, we diluted respective anti-stem Ab fourfold (starting at 100nM) in the presence or absence of C1q (220nM) in RPMI 1640 medium supplemented with 0.3% BSA. We added the mixture to IAV-infected MDC SIAT1 cells and continued as described above. We plotted the data using GraphPad Prism7 software.

### Protein gels, immunoblotting, and dot blotting

We diluted freshly purified virus samples in DPBS and combined three volumes of the virus with one volume of NuPAGE LDS Sample Buffer (4×; Thermo Fisher) containing 100 mM dithiothreitol. We heated samples at 70°C for 10 min, cooled them on ice, and loaded 10 µl (100 ng) of the sample on NuPAGE 4–12% Bis-Tris Protein Gel (Thermo Fisher), separating proteins by electrophoresis with Chameleon Pre-Stained Protein Ladder (Li-Cor) on 4–12% Bis-Tris Gels (Invitrogen) at 150 V for 90 min. We transferred proteins to nitrocellulose membranes using an iBlot device at the P3 setting for 7 min. We blocked membranes with OBB for one h at room temperature and incubated them with rabbit polyclonal anti-PR8 HA antibody (Sino Biological) (2,000× dilution) and CM1 mAb (1 µg/ml) diluted in OBB. We washed the membrane in DPBS containing 0.05% NP-40 three times. We incubated for one h at room temperature with IRDye 800CW goat anti-mouse or IRDye 680RD goat anti-rabbit secondary Ab (Li-Cor) diluted 10,000-fold in OBB. We washed the membrane three times with one additional Milli-Q water washing step. We scanned the membrane with the near-infrared imaging system (OdysseyCLx). We used ImageStudioLite software to quantify the signal and plotted the data using GraphPad Prism7 software. We scanned membranes with the Odyssey CLx scanner and quantified them with ImageStudio Lite software.

### Flow-cytometry-based IAV neutralization assay

We transferred 50,000 MDCK SIAT7e suspension cells diluted DPBS into polypropylene not treated round-bottom 96-well microplate (Corning). We preincubated (at 37°C for one h) 40 µl of half-log_10_ serially diluted Abs in Minimum Essential Media (MEM) media (Gibco) supplemented with 10mM Hepes (Gibco), 50 μg/ml Gentamicin (Gibco), 0.3% BSA (infection media) with 40 µl of the respective HA variant mCherry-expressing or regular (HA2: I6F) virus diluted in infection in polypropylene not treated round-bottom 96-well microplate. Alternatively, we added C1q at 160 nM into the infection medium. We spun MDCK S7e cells at 2,000 RPM for five minutes, aspirated PBS, and transferred 70 µl Ab-virus or C1q-Ab-virus mixture to washed MDCK S7e cells and incubated samples at 37°C for (six-seven h). We added paraformaldehyde (Electron Microscopy Science) to the final 1.6% for 20 minutes at four °C, and we added 100 ul DPBS. For the regular virus infection, we washed cells with DPBS, centrifuged cells at 2,000 RPM for five minutes, aspirated the supernatant, and fix-permeabilized the cells with a Foxp3/Transcription Factor Staining Buffer Set (Thermo Fisher) as recommended by the manufacturer. We washed the cells once with DPBS, incubated them with 1,000x diluted NP mAb (HB65-AF647) for 30 min, and washed and resuspended them with 0.1% BSA solution. We analyzed samples using a BD LSRFortessa X-20 instrument. Analysis was performed using FlowJo software (TreeStar). We normalized the frequencies of mCherry/NP-positive cells by the non-Ab–treated samples or plateaued response observed for at least two highest Ab dilutions employed (set to 100%) and fitted nonlinear regression curves using the dose-response inhibition model for neutralization assay with GraphPad Prism7 software.

### SARS-Covid2-EGFP and VSVΔG-EGFP-SARS2-Sgp pseudovirus neutralization assay

We transferred 50,000 freshly trypsinized BHK21-Ace2 cells diluted in DPBS into polypropylene not treated round-bottom 96-well microplate (Corning). For SARS-Covid2-EGFP neutralization, we followed the National Institute of Allergy and Infectious Diseases SARS-Cov-2 Virology Core SOPs. All steps involving the infectious SARS-Cov2-EGFP virus were performed at the BSL3 laboratory. We preincubated (at 37°C for one h) 40 µl of half-log_10_ serially diluted Abs in Minimum Essential Media (MEM) media (Gibco) supplemented with 10mM Hepes (Gibco), 50 ug/ml Gentamicin (Gibco), 0.3% BSA (infection media) with 40 µl of the virus diluted to MOI=0.01-0.015 in infection medium. Alternatively, we supplemented infection media with C1q (160nM). We spun BHK21-Ace2 at 2,000 RPM for five minutes and gently aspirated the supernatant. We transferred 70 μl of Ab-virus or C1q-Ab-virus mixture to cells and incubated cells overnight (12-15 h). We spun down the cells, aspirated the supernatant, and added 15 μl TrypLE Express (Gibco) for three minutes. We added 190 ul of 4% PFA (Electron Microscopy Science) resuspended cell and incubated for 30 minutes at room temperature. We transferred cell suspensions to a new plate, spun the cells down at 2,000 RPM for five minutes, and replaced PFA for DPBS supplemented with 0.3% BSA.

For the VSVΔG-EGFP-SARS2-Sgp pseudovirus neutralization assay, we repeated the protocol above except for the PFA inactivation step. We analyzed samples using a BD LSRFortessa X-20 instrument. Analysis was performed using FlowJo software (TreeStar). We normalized the frequencies of mCherry/NP-positive cells by the non-Ab–treated samples or plateaued response observed for at least two highest Ab dilutions employed (set to 100%) and fitted nonlinear regression curves using the dose-response inhibition model for neutralization assay with GraphPad Prism7 software.

### Biolayer interferometry

We performed human plasma and avidity variant Abs’ binding using an Octet Red384 biolayer interferometer (Sartorius). We completed all the steps in 0.05% BSA and 0.002% Tween20. We diluted human plasma samples tenfold. For the anti-stem Abs detection in human plasma samples, we used Dip and Read Octet® Anti-Penta-HIS (HIS1K) biosensors to load (spectral shift Δλ=0.35-0.5 nm) NC/99/H1N1 derived rHA-stem only protein (roughly 140 nM). We measured the association at 30°C for 1500 seconds (till saturation), followed by 1500 seconds of Fi6 (200 nM) binding competition.

For the binding characterization of the human IgG1 Ab to human and mouse C1q, respectively, we used FAB2G Dip and Read Octet Biosensors (Sartorius) specific for CH1 domain of Fab Ab’s portion. We loaded biosensors with the 310-16G8 IgG1-wt or – K322A (ΔC1q binding variant) till saturated (100 µg.ml^-1^, spectral shift Δλ=3.5-4 nm). We measured the association of the human (Complement Technology, Inc.) and mouse C1q (Complement Technology, Inc.) (100 µg.ml^-1^) at 30°C for 120 seconds. For competition, we saturated hIgG1-wt pre-loaded sensor with goat F(ab)_2_ anti-human-Fc (100 µg.ml^-1^, spectral shift Δλ=3.5-4 nm) and we measured subsequent association with human C1q. We plotted and analyzed data in GraphPad Prism 7.

### Animal intranasal passive transfer challenge studies

We purchased C57BL/6 mice from Taconic Farm. We used female 6-8-wk-old mice randomly assigned to experimental groups (n=six to seven). We held mice under specific pathogen–free conditions. All animal procedures were approved and performed in accordance with National Institute of Allergy and Infectious Diseases Animal Care and Use Committee Guidelines. For passive transfer, we administered each mAb (100 µg, 20 µg, 4 µg or 300ng per mouse 18-20 g) with or without human C1q (25 µg per mouse) in 25 µl sterile saline solution intranasaly under mild isoflurane anesthesia (3.5 %). As a negative control we treated mice with C1q only. One hour later, we anesthetized mice with isoflurane as previously described and administered 25 µl virus intranasally (PR8, 3 × 10^3^ TCID_50_) diluted in sterile PBS. We recorded weight for 14 d and euthanized mice when mice reached 30% weight loss (an experimental endpoint).

## Acknowledgments

This work was supported by the Division of Intramural Research of the National Institute of Allergy and Infectious Diseases. We thank the Department of Transfusion Medicine (NIH/CC/DTM) for providing human blood-derived products.

We thank Patrick C. Wilson (Anne E. Dyson Professor of Pediatric Research for providing Abs–expressing plasmids. We are grateful to Bernard Lafont, Johnson Reed, and Nicole Lackemeyer, the NIAID SARS-CoV-2 Virology Core BSL3 Laboratory staff, for their excellent support, training, and the help provided.

We thank Jojeph Shiloach (NIH/NIDDK) for providing suspension MDCK SIAT7e cell line.

The authors declare no competing financial interests.

## Supplementary Material

**Supplemental Table 1.** Characteristics of HA mAbs used.

| Ag site | Sa |  | Sb |  |  |  |  | Ca |  |  | Cb |  |  | stem |  |  |  |  |  |
| --- | --- | --- | --- | --- | --- | --- | --- | --- | --- | --- | --- | --- | --- | --- | --- | --- | --- | --- | --- |
| Ab | Y8-1A6 | H2-6A1 | H36-26 | H36-4 | H28-E23 | IC5-4F8 | H28-D14 | H17-L10 | H17-L2 | H2-4B1 | H9-D3 | H35-C12 | H17-L7 | 310-16G8 | 310-18F8 | 310-18E7 | Fi6 | 310-18D5 | CR9114 |
| K<sub>D</sub> nM | 1.3 | 1.0 | 0.1 | 0.2 | 0.1 | 0.1 | 0.1 | 3.1 | 2.4 | 17.1 | 1.5 | 2.4 | 0.7 | 0.39 | 0.8 | 0.9 | 0.5 | 0.3 | 1.6 |
| Isotype | mlgG2a | mlgG2a | mlgG2a | mlgG2a | mlgG2a | mlgG3 | hlgG1 | mlgG2a | mlgG2a | mlgG1 | mlgG3 | hlgG1 | mlgG2a | hlgG1 | hlgG1 | hlgG1 | hlgG1 | hlgG1 | hlgG1 |

**Supplemental Table 2.** SARS-Cov-2 mAbs used.

| Ab | domain | reference |
| --- | --- | --- |
| COVA1-18 | RBD | Brouwer<sup>59</sup> |
| COVA2-39 | RBD | Brouwer<sup>59</sup> |
| COV2-2832 | RBD | Zost<sup>58</sup> |
| COV2-2807 | RBD | Zost<sup>58</sup> |
| COV2-2780 | RBD | Zost<sup>58</sup> |
| c102 | RBD | Barnes<sup>60</sup> |
| c119 | RBD | Barnes<sup>60</sup> |
| C105 | RBD | Barnes<sup>60</sup> |
| c110 | RBD | Barnes<sup>60</sup> |
| c121 | RBD | Barnes<sup>60</sup> |
| c135 | RBD | Barnes<sup>60</sup> |
| COVA2-42 | nonRBD | Brouwer<sup>59</sup> |
| COVA1-25 | nonRBD | Brouwer<sup>59</sup> |
| COVA1-02 | nonRBD | Brouwer<sup>59</sup> |
| COVA1-06 | nonRBD | Brouwer<sup>59</sup> |
| COVA1-09 | nonRBD | Brouwer<sup>59</sup> |
| COVA2-10 | nonRBD | Brouwer<sup>59</sup> |
| COVA2-30 | nonRBD | Brouwer<sup>59</sup> |
| COVA1-12 | nonRBD | Brouwer<sup>59</sup> |
| COVA2-28 | nonRBD | Brouwer<sup>59</sup> |
| COVA2-33 | nonRBD | Brouwer<sup>59</sup> |
| COVA1-22 | NTD | Brouwer<sup>59</sup> |

**Supplemental Table 3.**
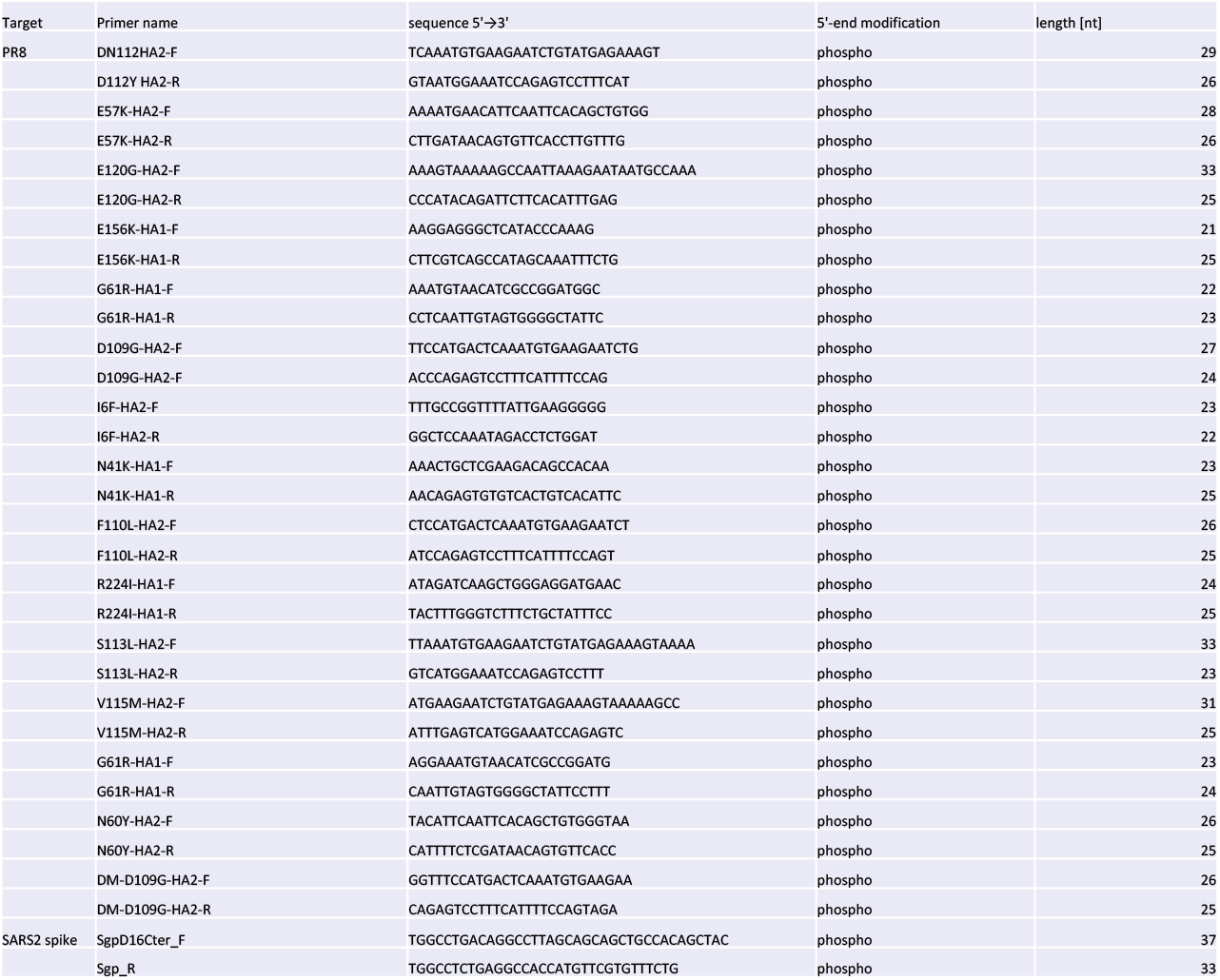
Primers.

**Supplemental Figure 1.**
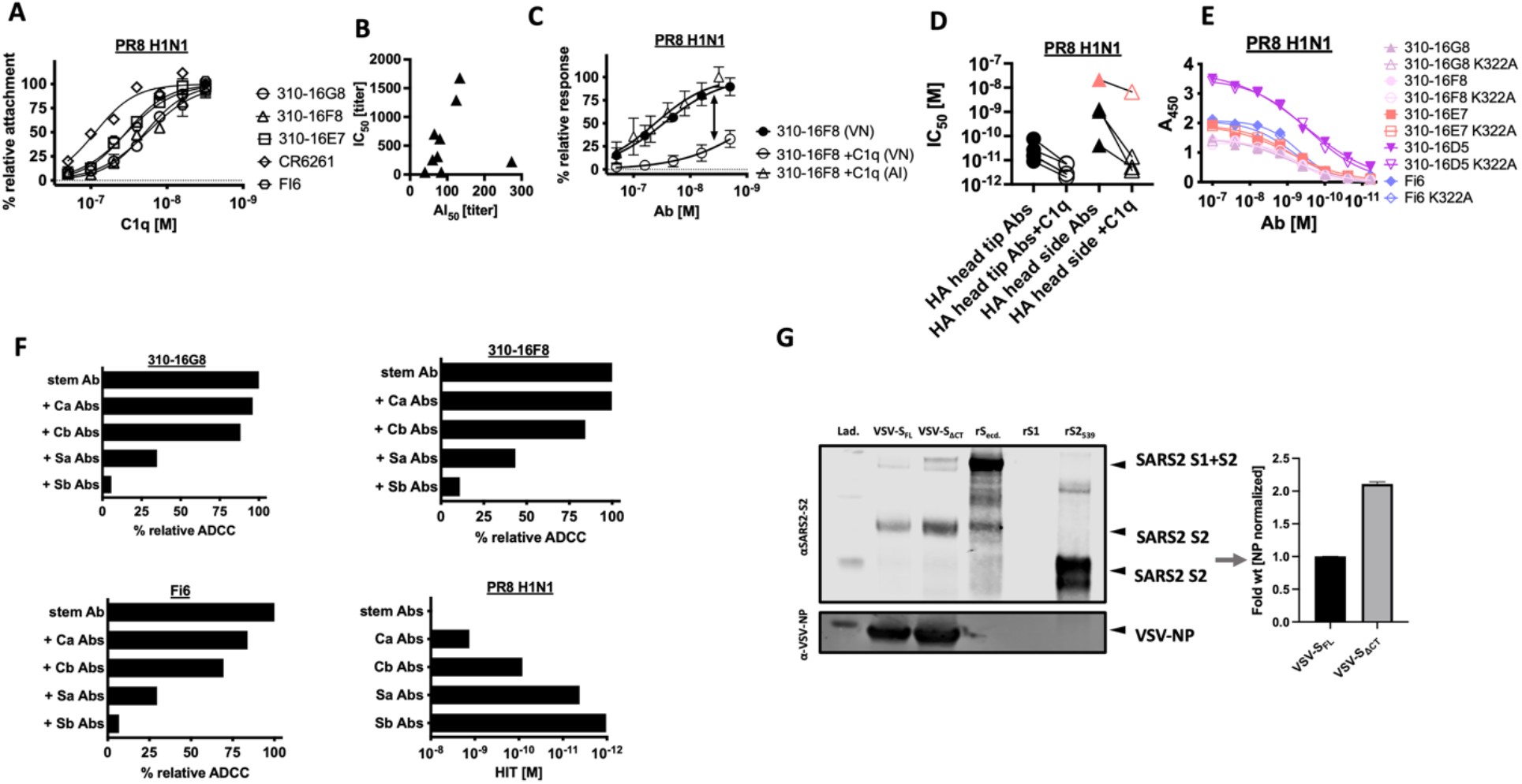
**(A)** AI activity of the indicated anti-stem and head mAbs (constant 100nM) in the presence of graded C1q *vs.* PR8 determined using the fetuin-attachment assay. **(B)** IC_50_ vs AI_50_ for respective human plasma samples. **(C)** C1q-AI, C1q-VN, and VN of the indicated mAb vs. PR8 in the fetuin-attachment assay in single cycle flow neutralization assay. Arrow indicates concentration with negligible AI and VN unless C1q is present. **(D)** IC_50_ of indicated HA head mAbs vs PR8 in the presence or absence of C1q in the single cycle flow neutralization assay. **(E)** ELISA-determined anti-stem mAb wt vs. K322A binding to PR8. **(F)** ADCC of stem mAbs indicated (alone or mean response determined for stem mAb combined with individual Ca, Cb, Sa, Sb specific mAbs) vs PR8 infected MDCK Siat1 cells in Raji-NFAT-Lucifarase ADCC reporter assay. Bottom right is HI of respective antigenic site specific mAbs vs PR8. **(G)** Relative amount of wt and ΔCT SARS-Cov2 Spike protein incorporated into VSV pseudo viral particles in WB. A mean of four to six replicates acquired in two to three independent is presented. Error bars indicate the SD. Non-primary antibody control was included in ELISA. The NVC and NoAbC were included in each experiment. Data are normalized by NoAbC and/or by plateaued response observed for at least two highest Ab dilutions employed (set to 100%). For ADCC stem mAb response alone set to 100%. Data are plotted and analyzed with GraphPad Prism7 software. The nonlinear regression curves were fitted using the specific binding with a hill slope regression for the ELISA, the dose-response inhibition model was used for the VN and AI assays, and IC_50_ values were calculated.

**Supplemental Figure 2.**
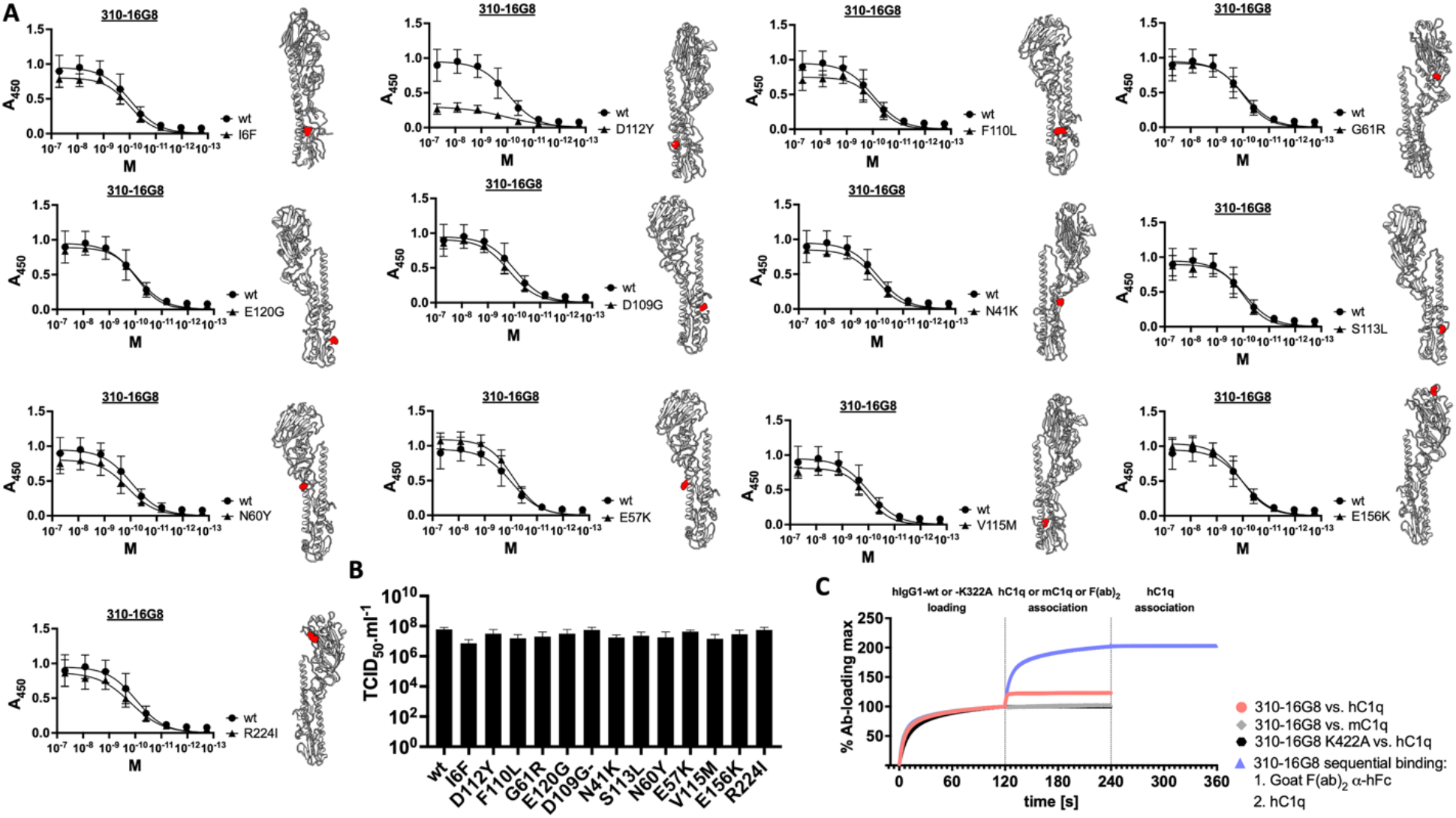
**(A)** ELISA-determined anti-stem mAb binding to PR8 vs escape variangs indicated. Locations of amino acid substitutions on HA monomers are shown in red. Each experiment shows the mean of four technical replicates acquired in two independent experiments. Error bars indicate the SD. The non-primary antibody control was included. We Data vere plotted and analyzed with GraphPad Prism7 software. The nonlinear regression curves were fitted using the specific binding with a hill slope for ELISA. **(B)** We determined endpoint viral growth titers *in ovo*. **(C)** Binding of human IgG1-wt vs. -K322A to human and mouse C1q respectively.

